# Human cellular model systems of β-thalassemia enable in-depth analysis of disease phenotype

**DOI:** 10.1101/2022.09.01.506225

**Authors:** Deborah E Daniels, Ivan Ferrer-Vicens, J Hawksworth, Tatyana N Andrienko, Elizabeth M Finnie, Daniel C J Ferguson, A. Sofia F. Oliveira, Jenn-Yeu A. Szeto, Marieangela C Wilson, Jan Frayne

## Abstract

β-thalassemia is a prevalent genetic disorder causing severe anemia due to defective erythropoiesis, with few treatment options. Studying the underlying molecular defects is impeded by paucity of suitable patient material. In this study we created human disease cellular model systems for β-thalassemia, which accurately recapitulate the phenotype of patient erythroid cells. We also developed a high throughput compatible fluorometric-based assay for evaluating severity of disease phenotype and utilised the assay to demonstrate positive response of lines to verified reagents, providing validation for such applications.

TMT-based comparative proteomics confirmed the same profile of proteins previously reported, whilst providing new insights into the altered molecular mechanisms in β-thalassemia erythroid cells, with upregulation of a wide range of biological pathways and processes observed.

Overall, the lines provide a sustainable supply of disease cells as novel research tools, for identifying new therapeutic targets, and as screening platforms for novel drugs and therapeutic reagents.

## Introduction

β-thalassemia syndromes are a heterogeneous range of anemias and a major source of morbidity, mortality, and substantial financial burden globally. The disease is characterised by variable reduced (β^+^) or absent (β^0^) β-globin chain synthesis, with more than 300 different mutations in and around the β globin gene identified^1,2^. Mutations resulting in β^0^-thalassemia cause a severe chronic anemia, whereas that resulting from β^+^ varies from clinically benign to severe.

Reduction in β-globin results in excess α-globin, forming insoluble aggregates which undergo auto-oxidation leading to increased reactive oxygen species (ROS) and a range of intracellular downstream events that result in ineffective erythropoiesis (IE), the hallmark and primary cause of β-thalassemia disease pathophysiology. IE is manifest as increased expansion of erythroid progenitors due to increased EPO production *in vivo*, but with a maturation block at the polychromatic stage of differentiation^3–5^ with increased apoptosis (reviewed by Ribeil et al.^6^), and thus impaired erythropoiesis. However, the underlying molecular mechanisms that cause IE are not fully understood.

In recent years there has been much renewed interest in studying β-thalassemia, however there are still few treatment options. The mainstay is blood transfusion, but this has associated complications of alloimmunisation and iron-mediated potentially fatal organ damage, requiring life-long iron-chelation therapy.

Non-curative approaches have aimed to correct globin imbalance by promoting γ -globin synthesis, such as with hydroxyurea^7^, but responses are inconsistent^8–10^. More recently, luspatercept has been shown to increase hemoglobin values, but modestly by 1-2 g/dl in non-transfusion dependent thalassemia patients (NTDT). However, only 20% of transfusion-dependent (TDT) patients showed a reduction of blood transfusion by more than a third^11^.

The only definitive cure is bone marrow transplant, applicable for some young patients with a matched sibling lacking TDT. Gene therapy, either using gene replacement or editing with associated bone marrow transplantation hold much promise for those lacking a matched donor. However, efficacy and safety issues are still concerns and given the complexity and high cost of these approaches, they are unlikely to scale to the massive global unmet need for β-thalassemia therapeutics, particularly in low to middle income countries. Thus, new cost-effective treatment strategies and drug targets are desperately required to deliver optimal therapies to the greatest numbers of people.

However, studying the molecular defects underlying the β-thalassemia phenotype, and identifying drug targets, is severely impeded by paucity of suitable material from patients, and lack of suitable cell lines. Although erythroid cells can be generated *in vitro* from peripheral blood stem cells, the approach is severely limited by the restricted expansion potential of the cells^12^ and thus number of erythroid cells generated, with repeat collections required, a particularly unsuitable approach for anemic patients. Thus, currently much of our understanding is based on mouse models which are also used to investigate and evaluate the effect of drugs. However, fundamental differences between mouse and human erythropoiesis are becoming increasingly apparent^13,14^. Therefore, new cellular models for β-thalassemia are essential to enable in-depth investigation of underlying molecular mechanisms, to aid identification of new therapeutic targets and as screening platforms for the effect and efficacy of potential new drugs and reagents.

In this study we created and validated novel human disease erythroid cell lines for β-thalassemia. Homozygous (*HBB*^−/-^) and heterozygous (*HBB*^+/-^) *HBB* gene knockout mutations, and the prevalent β-thalassemia mutations CD41/42 -TTCT and IVS-1-1 G→ T, were introduced into the erythroid cell line BEL-A, which undergoes normal erythropoiesis and expresses a normal adult globin profile, via CRISPR genome editing. All disease lines accurately recapitulate the disease phenotype of β^0^-thalassemia patient erythroid cells, whilst the *HBB*^+/-^ lines recapitulate cells of individuals with β-thalassemia trait. The lines thus provide a sustainable and consistent supply of disease cells as unique research tools, and as drug screening platforms. For the latter we also developed a high throughput compatible flow cytometry assay for evaluation of severity of IE. We utilised the assay to demonstrate that the *HBB*^−/-^ line responds appropriately to both hydroxyurea treatment and *BCL11A* +58 enhancer editing, with increased HbF and correlated improvement to IE, validating the line for screening applications.

We also performed TMT-based comparative proteomic analysis. This confirmed the same profile of proteins previously reported dysregulated in β-thalassemia erythroid cells, providing further validation, whilst also providing novel data on the altered molecular mechanisms in β-thalassemia erythroid cells. This included increased levels of antioxidant response enzymes, heat shock proteins and chaperones, and many proteins of the ubiquitin-proteasome and autophagy pathways along with proteins associated with lysosome biogenesis, together with up-regulation of respective pathways. The data also revealed upregulated pathways and increased levels of proteins for heme biosynthesis, along with associated upregulated endosome biogenesis and trafficking pathways, which may contribute to the disease phenotype and provide potential new therapeutic targeting opportunities.

Finally, we also describe a novel β-globin splice variant arising from the IVS-1-1 G→ T mutation which is unable to form stable tetramers with α-globin, the variant instead precipitating and potentially contributing to the IE phenotype in IVS-1-1 erythroid cells.

## Results

The erythroid cell line BEL-A recapitulates normal adult erythroid cell differentiation and has a normal adult globin expression profile^15^. It is therefore an excellent founder line to introduce mutations to create cellular model systems of red blood cell diseases, and so was used in this study to create β-thalassemia disease lines.

### Generating HBB^+/-^ and HBB^−/-^ erythroid progenitor cell lines

As β-thalassemia is caused by a wide range of mutations that inhibit or reduce the production of β-globin, in the first instance homozygous (*HBB*^−/-^) and control heterozygous (*HBB*^+/-^) β-globin knockout lines were created using CRISPR-Cas9 mediated genome editing of BEL-A. This resulted in biallelic 1 base pair and monoallelic 2 base pair deletions for the *HBB*^−/-^ and *HBB*^+/-^ clonal lines respectively (Fig. 1A), leading to frameshift and premature stop codons in exon 1 in both cases.

**Figure 1:**
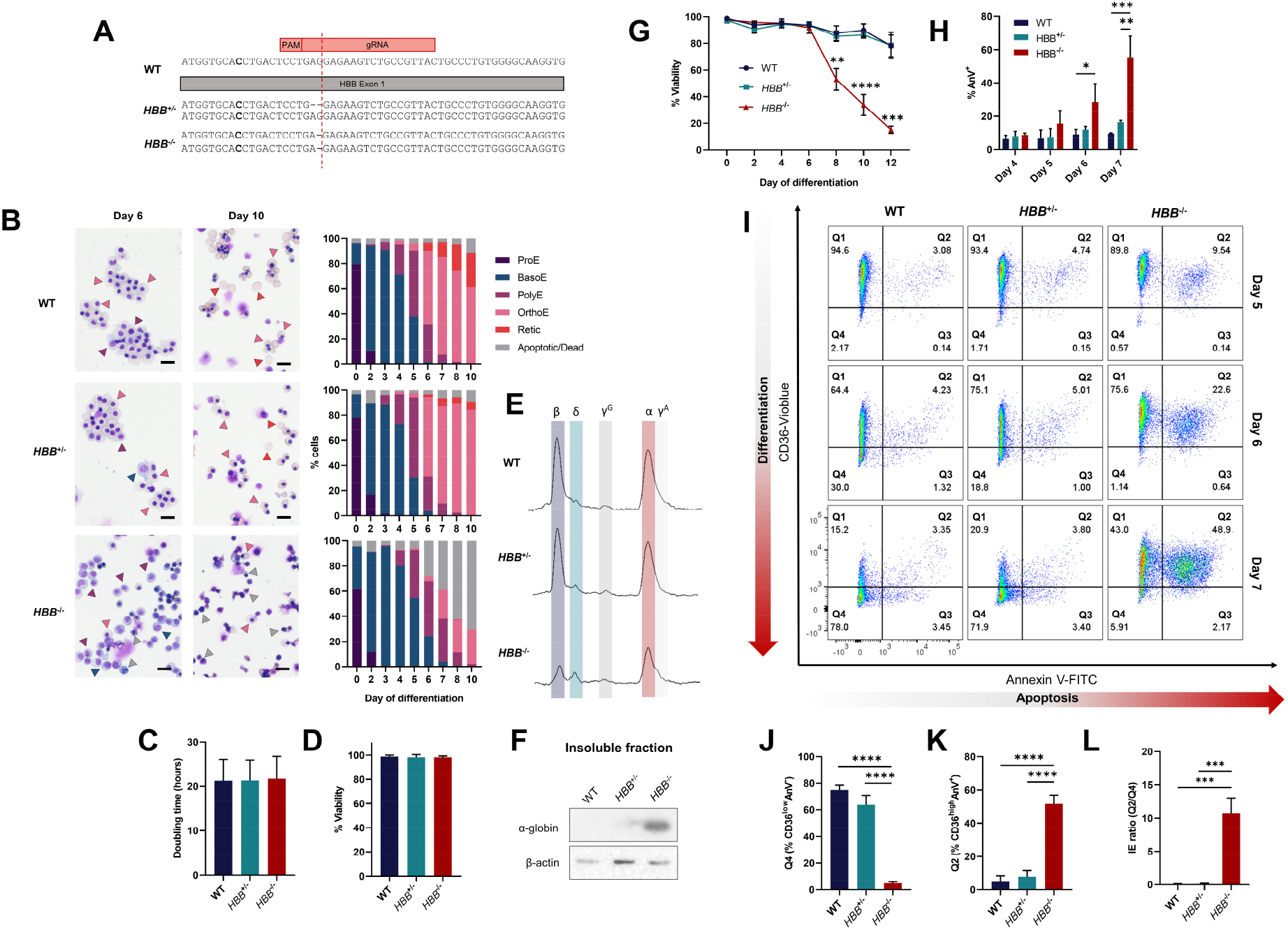
Generation and characterisation of *HBB*^+/-^ and *HBB*^−/-^ erythroid progenitor cell lines. (A) Schematic of DNA sequence for *HBB*^+/-^ and *HBB*^−/-^ compared to WT BEL-A cells. (B) Representative cytospins of WT, *HBB*^+/-^ and *HBB*^−/-^ BEL-A with morphology quantification (ProE, proerythroblast; BasoE, basophilic erythroblast; PolyE, polychromatic erythroblast; OrthoE, orthochromatic erythroblast; Retic, reticulocyte). Arrowheads indicate the following cell types: blue, BasoE; purple, PolyE; pink, OrthoE; red, Retic; grey, dead/apoptotic. Scale bars 20 μm. Doubling time (C) and % viability (D) of expanding cell lines as determined by trypan blue exclusion assay, n=14. (E) RP-HPLC traces for WT, *HBB*^−/-^ and *HBB*^+/-^ BEL-A at day 6 of differentiation. Peaks are identified for β-globin (β), δ-globin (δ), ^G^γ-globin (^G^γ), α-globin (α) and ^A^γ-globin (^A^γ). (F) Representative western blot of aggregate proteins from WT, *HBB*^+/-^ and *HBB*^−/-^ BEL-A harvested at day 6 of differentiation, incubated with α-globin antibody. β-actin was used as a protein loading control. (G) Percentage viability of WT, *HBB*^−/-^ and *HBB*^+/-^ BEL-A during differentiation by trypan blue exclusion assay. (H) Percentage of Annexin V positive cells (AnV^+^) in WT, *HBB*^+/-^ and *HBB*^−/-^ BEL-A during differentiation as determined by flow cytometry. (I) Representative IE flow cytometry plots at day 5, 6 and 7 of differentiation. Quantification of Q4 CD36^low^AnV^−^ cells (J), Q2 CD36^high^AnV^+^ (K) and IE ratio (Q2/Q4) (L) at day 7 of differentiation. Results show mean ± SD, n=3 unless otherwise stated. \**P* < .05, \*\**P* < .01, \*\*\**P* < .001, \*\*\*\**P* < .0001.

The resultant *HBB*^+/-^ and *HBB*^−/-^ lines had a predominantly pro-erythroblast morphology with some basophilic erythroblasts, in line with the WT BEL-A line (Fig. 1B). Doubling times and viability were invariant across the lines (Fig. 1C and D).

Analysis of globin levels by RP-HPLC showed that *HBB*^*+/-*^ cells produced similar levels of all globin subunits to WT cells whereas, as expected, β-globin was eliminated in the *HBB*^−/-^ line (Fig. 1E; see also Supplementary Fig. 1 for clarification of residual peak overlapping position of β-globin). α -globin was also reduced in the *HBB*^−/-^ cells as, in absence of β-(or γ -) globin it forms insoluble aggregates (Fig. 1F), a feature of β-thalassemia patient erythroid cells.

When differentiated, viability of the WT and *HBB*^+/-^ line remained high throughout the culture duration (Fig. 1G). In contrast the *HBB*^−/-^ line recapitulated the phenotype of β 0-thalassemia erythroid cells^3^ with clear ineffective erythropoiesis, as demonstrated by a dramatic and significant decline in viability from day 6 of differentiation (Fig. 1G), corresponding temporally with differentiation to polychromatic erythroblasts (Fig. 1B), and with a significant increase in Annexin V (AnV) cell surface abundance (Fig. 1H), indicative of apoptosis (Fig. 1B). In line, growth curves during differentiation showed unchanged expansion in HBB^+/-^ and HBB^−/-^ lines compared to WT up until day 6, but with greater reduction in cell numbers post-day 6 in HBB^−/-^ line due to cell death (Supplementary Fig. 2). Similarly, kinetics of the *HBB*^+/-^ cultures were invariant to that of WT cells during differentiation (Fig. 1B). In contrast, although the majority of *HBB*^*-/-*^ cells differentiated to polychromatic erythroblasts, few differentiated further to orthochromatic erythroblasts (Fig 1B), as found for cultured patient erythroid cells^3^.

Analysis of a second *HBB*^−/-^ clonal line (*HBB*^−/-^C2) displayed the same phenotype, with similar striking decrease in viability from day 6 of culture and associated significant increase in AnV levels and apoptosis at the polychromatic stage (Supplementary Fig. 3A-C).

Overall, the data confirm the *HBB*^−/-^ lines as robust cellular models of β^0^-thalassemia, providing a sustainable and consistent supply of disease cells.

### Assay for assessing severity of ineffective erythropoiesis

To evaluate drugs and reagents that reduce severity of IE, thus improving disease phenotype, the WT, *HBB*^+/-^ and *HBB*^−/-^ lines were used to establish a flow cytometric assay for IE utilising fluorophore tagged antibodies to membrane markers. For this, the abundance of membrane protein CD36, which decreases during erythroid cell differentiation^16,17^, was first evaluated. CD36 levels decreased from day 6 of differentiation in WT and *HBB*^+/-^ cultures, but was retained in *HBB*^−/-^ cultures, correlating with arrested differentiation of *HBB*^−/-^ cells at the polychromatic stage (Supplementary Fig. 4), and providing effective resolution of transition from polychromatic to orthochromatic erythroblasts. Dual staining of CD36 with Annexin V then provided a robust and high throughput compatible assay for monitoring IE, as illustrated in Figure 1I. To clarify, as differentiation proceeds WT and *HBB*^+/-^ cells show progressively decreasing levels of CD36 and consistently low levels of AnV, with the majority of cells by day 7 in quadrant 4, which delineates viable orthochromatic erythroblasts (CD36^low^AnV^−^; Fig 1I and J). In contrast very few *HBB*^−/-^ cells proceed to quadrant 4, instead shifting from quadrant 1 to 2 (CD36^high^AnV^+^; Fig 1I and K) as levels of apoptosis increase with cells reaching the polychromatic stage. The number of cells in Q2 (CD36^high^AnV^+^):Q4 (CD36^low^AnV^−^) provides an IE ratio and numerical value for the severity of IE (Figure 1L). The second *HBB*^−/-^ clone showed similar profile and IE ratio in the assay (Supplementary Fig. 3D-E).

### Validating β^0^-thalassemia line as a drug and reagent screening platform

To test the potential of the *HBB*^−/-^ line as a screening platform for novel drugs and therapeutic approaches for reducing the disease severity of β-thalassaemia, we employed two established approaches that increase the level of γ-globin, and thus fetal hemoglobin (HbF) in erythroid cells. Upregulation of γ-globin was selected as our gauge, as it is the most promising approach for the treatment of β -thalassemia, and also sickle cell disease (SCD), compensating for loss of, or mutated, β -globin.

Hydroxyurea (HU) increases γ-globin expression in human primary erythroid cells^18^ and is currently approved for the treatment of non-transfusion dependent β-thalassemia and SCD. Consistent with previous findings, differentiated *HBB*^−/-^ cells showed a dose-dependent increase in the proportion of HbF^high^ cells (Fig. 2A and C) and total γ-globin level by RP-HPLC (Supplementary Fig. 5) in response to HU treatment. To equate with effect on disease severity, cells were evaluated by our IE assay, which revealed a concurrent decrease in IE ratio with increasing concentrations of HU (Fig. 2B and D). We also used CRISPR-Cas9 gene editing to disrupt the +58 enhancer of the γ-globin repressor *BCL11A* in *HBB*^−/-^ cells, using a previously validated sgRNA^19,20^, an approach granted PRIME designation for TDT and SCD, with planned regulatory submission this year. A clonal population of +58 enhancer edited *HBB*^−/-^ cells containing a compound heterozygous deletion (Supplementary Fig. 6A) showed an ∼25% increase in γ-globin levels (Supplementary Fig. 6B and C) in line with previous findings^21^, and a decrease in the severity of IE, as indicated by an ∼50% decrease in the IE ratio at day 7 of differentiation and an increase in cell viability at day 8 and 10 of culture (Supplementary Fig. 6D-F).

**Figure 2:**
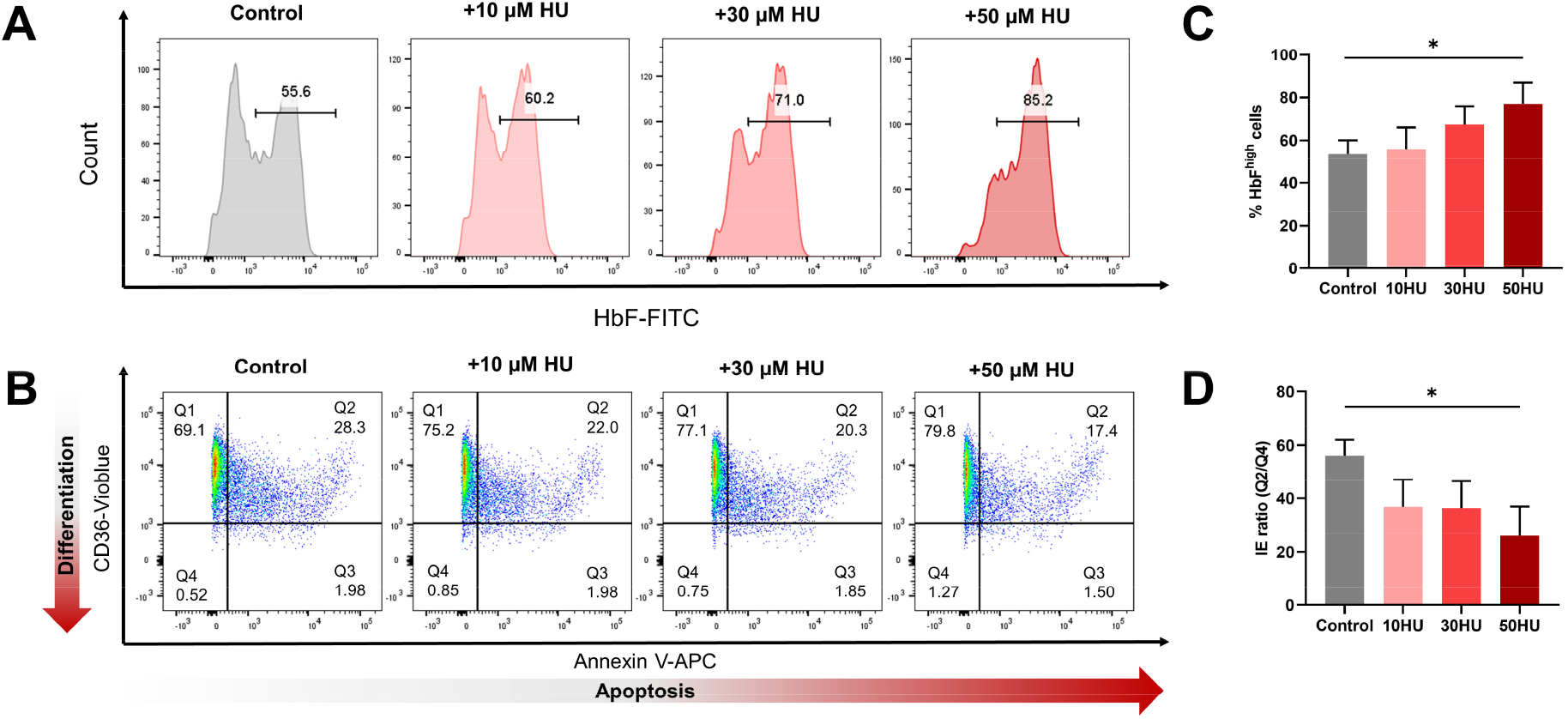
*HBB*^−/-^ BEL-A responds to hydroxyurea (HU) to increase γ-globin expression and reduce IE. Representative flow cytometry plots of control and hydroxyurea (HU) treated *HBB*^−/-^ BEL-A cells at day 7 of differentiation showing distribution of HbF-FITC abundance (A) and IE measured by CD36-vioblue vs Annexin V-APC (B). (C) Quantification of HbF^high^ cells. (D) Quantification of IE ratio (%CD36^high^AnV^+^/ CD36^low^AnV^−^). Results show mean ± SD, n=3. \**P* < .05.

These findings confirm that the *HBB*^−/-^ disease line responds effectively to both pharmacological and genome editing approaches to increase γ-globin expression, whilst also directly demonstrating that the treatments reduce IE and thus the severity of β^0^-thalassemia disease phenotype, collectively providing proof-of-principle for utilising the line for screening applications.

### Generating β^0^-thalassemia cell lines with specific prevalent mutations

While the *HBB*^−/-^ lines do successfully recapitulate the expected phenotype of β^0^-thalassemia erythroid cells, the deletion mutation is not one of over 300 different mutations that cause reduced or absent β-globin production in β-thalassemia patients^1^. Therefore, we next selected two prevalent β^0^-thalassemia mutations to introduce into BEL-A, to more accurately replicate patient genotypes and corroborate data from *HBB*^−/-^ lines; CD41/42 -TTCT deletion which results in a frameshift and introduction of a premature stop codon^22,23^ and IVS-1-1 G→ T substitution which prevents correct splicing of *HBB* mRNA^23^.

Both homozygous mutations were successfully introduced into BEL-A using RNP-based CRISPR-Cas9 genome editing with a single-strand oligodeoxynucleotide (ssODN) donor template (Fig. 3A). In the case of IVS-1-1, due to the relatively large 24 base pair distance from the cleavage site of the closest possible *HBB*-specific sgRNA and the intended edit, the ssODN was recoded with silent mutations, which has been shown to enhance incorporation efficiency of distal edits^24^.

As for the WT line, both disease lines had a predominantly pro-erythroblast morphology with some basophilic erythroblasts (Fig. 3B).

**Figure 3:**
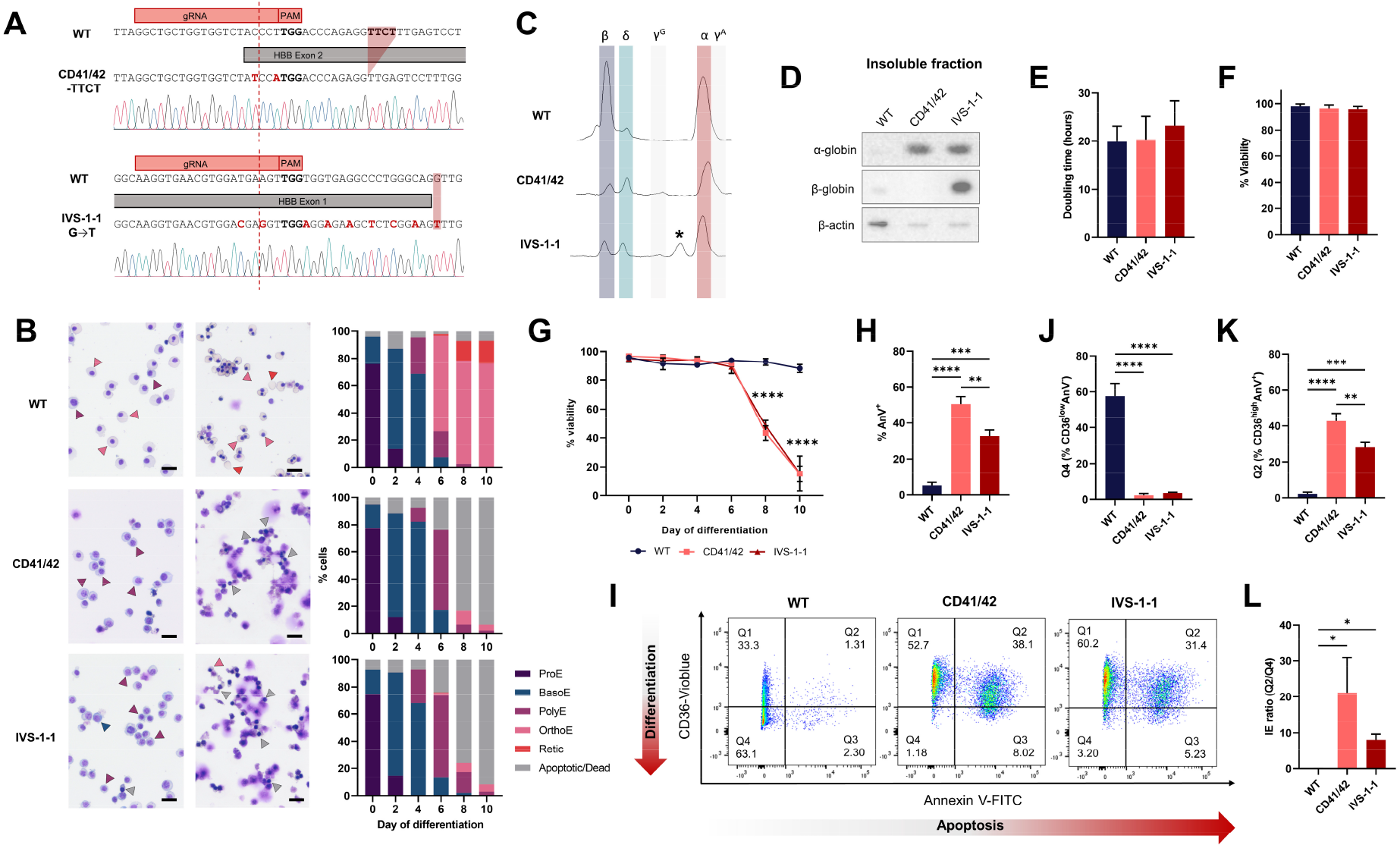
Generation and characterisation β^0^-thalassemia cell lines with specific prevalent mutations. (A) Schematic of DNA sequence for CD41/42 -TTCT and IVS-1-1 G→ T compared to WT BEL-A cells. (B) Representative cytospins of WT, CD41/42, and IVS-1-1 BEL-A with morphology quantification (ProE, proerythroblast; BasoE, basophilic erythroblast; PolyE, polychromatic erythroblast; OrthoE, orthochromatic erythroblast; Retic, reticulocyte). Arrowheads indicate the following cell types: blue, BasoE; purple, PolyE; pink, OrthoE; red, Retic; grey, dead/apoptotic. Scale bars 20 μm. (C) RP-HPLC traces for WT, CD41/42, and IVS-1-1 BEL-A at day 6 of differentiation. Peaks are identified for β-globin (β), δ-globin (δ), ^G^γ-globin (^G^γ), α-globin (α) and ^A^γ-globin (^A^γ). * Denotes splice β-globin variant (see Supplementary Figure 7). (D) Representative western blot of aggregate proteins from WT, CD41/42, and IVS-1-1 BEL-A harvested at day 6 of differentiation, incubated with α-globin or β-globin antibody. β-actin was used as a protein loading control. Doubling time (E) and % viability (F) of expanding cell lines as determined by trypan blue exclusion assay, n= 32. (G) Percentage viability of WT, CD41/42, and IVS-1-1 BEL-A during differentiation by trypan blue exclusion assay. (H) Percentage of Annexin V positive (AnV^+^) cells in WT, CD41/42, and IVS-1-1 BEL-A at day 7 of differentiation as determined by flow cytometry. (I) Representative IE flow cytometry plots at day 7 of differentiation. Quantification Q4 CD36^low^AnV^−^ cells (J), Q2 CD36^high^AnV^+^ (K) and IE ratio (Q2/Q4) (L) at day 7 of differentiation. Results show mean ± SD, n=3 unless otherwise stated. \**P* < .05, \*\**P* < .01, \*\*\**P* < .001, \*\*\*\**P* < .0001.

Similar to the *HBB*^−/-^ line, β-globin protein was eliminated in both the CD41/42 and IVS-1-1 clonal lines (Fig. 3C), and α -globin reduced due to insoluble aggregation (Fig. 3C and D), with doubling times and viability during the expansion phase not significantly different (Fig. 3E and F).

As for the *HBB*^−/-^ lines, both the CD41/42 and IVS-1-1 lines demonstrated IE, with significantly reduced cell viability at the polychromatic cell stage (from day 6; Fig. 3B and G), due to significantly increased apoptosis compared to control cells (Fig. 3H), with very few cells surviving and differentiating to orthochromatic erythroblasts (Fig 3G). Analysis using the flow cytometry IE assay showed similar profiles to the *HBB*^−/-^ line, with very few CD41/42 and IVS-1-1 cells shifted to quadrant 4 by day 7 (CD36^low^AnV^−^; Fig. 3I and J), instead having shifted to quadrant 2 (CD36^high^AnV^+^; Fig. 3I and K), due to levels of apoptosis increasing when cells reached the polychromatic stage, giving significantly increased IE ratios compared to WT cells (Fig. 3L).

Interestingly, an additional peak was observed in the IVS-1-1 line RP-HPLC traces (indicated by * in Fig. 3B). Mass spectrometry sequencing revealed this to be a splice variant of β-globin, created by splicing machinery using a previously identified IVS-1-13 alternative splice site^25^, resulting in a 4 amino acid insertion in β-globin between exon 1 and 2 (G29_R30insSLVS; Fig. 4A and Supplementary Fig. 7). Molecular modelling revealed disruption of interactions between the β-globin variant and α-globin, including with α-globin Phe 117, critical for α1β1 dimerisation^26^, due to increased distances between the network of key residues (Fig. 4B). Of note, the variant β-globin was detected in the insoluble fraction of IVS-1-1 cells (Fig. 3C), further supporting the hypothesis that it is unable to form a stable hemoglobin tetramer with α-globin and thus precipitating, potentially contributing to the IE phenotype of β-thalassemia erythroid cells with the IVS-1-1 mutation.

**Figure 4:**
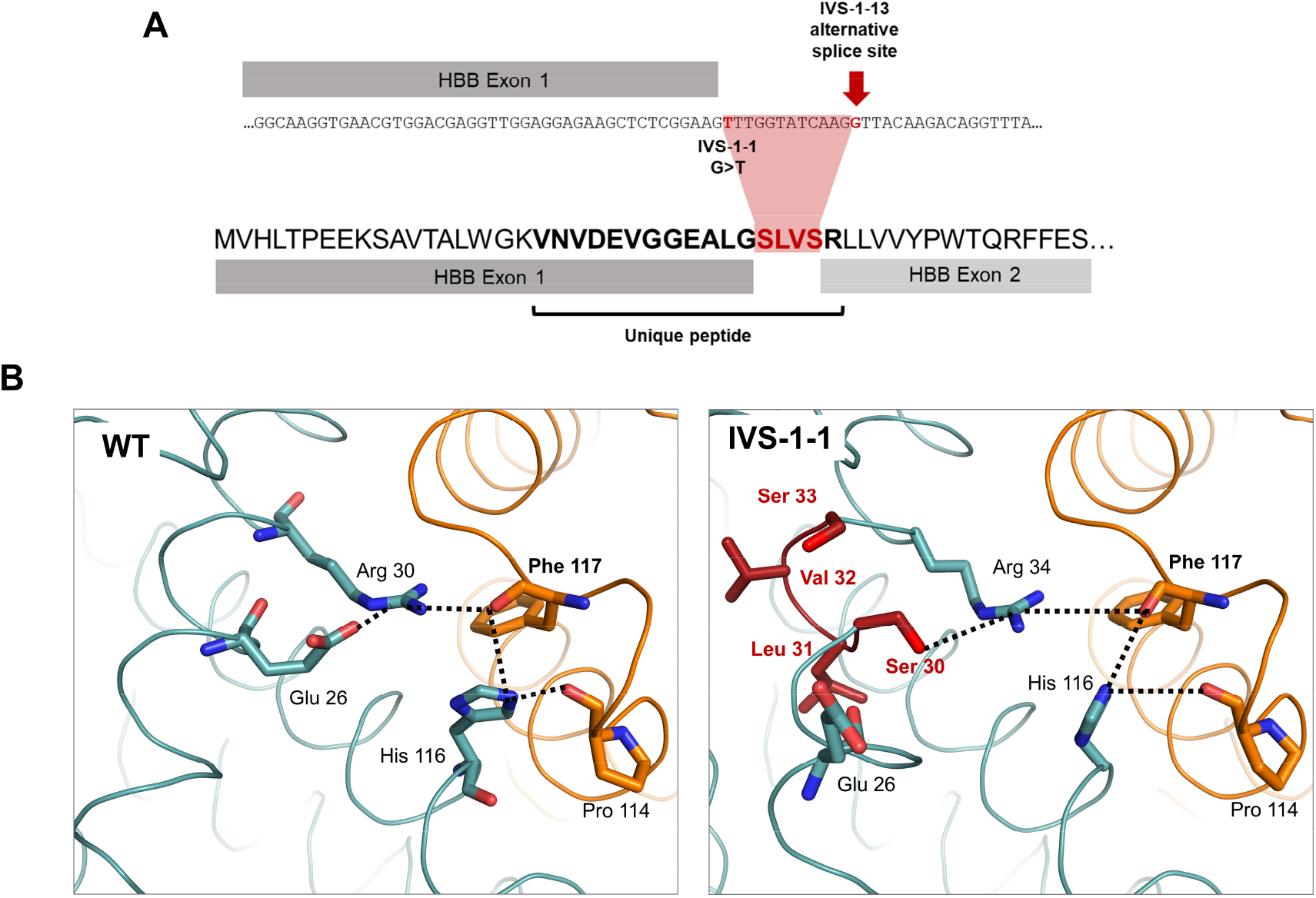
Identification of IVS-1-1 β-globin variant. (A) Schematic of the IVS-1-1 variant amino acid sequence resulting from use of the IVS-1-13 alternative splice site. The resulting peptide sequence contains a 4 amino acid insertion (red) from 12 base pairs of translated *HBB* intron. (B) Models of the α-/β-globin interface (α-globin in orange and β-globin in teal) showing increased distances between residues contributing to interactions with Phe 117 (dashed lines) on the α-globin chain in IVS1-1 compared to WT dimers. Inserted amino acids in the IVS-1-1 variant are shown in dark red.

### Quantitative proteomic comparison of WT and HBB^−/-^ BEL-A

To obtain a global assessment of differences between WT and *HBB*^−/-^ erythroid cell proteome, verify alterations in proteins previously identified, and reveal novel proteins and processes for potential therapeutic targeting, TMT LC-MS/MS comparative proteomic analysis was performed. For this, cells were evaluated during early differentiation (basophilic erythroblasts), prior to manifestation of IE phenotype and in mid-differentiation (polychromatic erythroblasts), when the α-/β-globin chain imbalance approaches maximal levels and apoptosis is induced. Due to differences in kinetics of the WT and *HBB*^−/-^ cultures, basophilic and polychromatic erythroblasts were isolated by FACS, using inverse levels of GPA and CD36 (representative cytospin images of these populations are shown in Supplementary Fig. 8).

Of the 3364 unique proteins quantified, 1048 (31%) were significantly different in *HBB*^−/-^ compared to WT basophilic erythroblasts, and 1001 (30%) in *HBB*^−/-^ compared to WT polychromatic erythroblasts (Table 1). Despite the similar proportion of significantly different proteins in both cell types, principal component analysis (PCA; Fig. 5A) illustrates the increased magnitude of proteomic changes between WT and *HBB*^−/-^ polychromatic erythroblasts compared to basophilic erythroblasts, reflecting the more severe phenotype of the former.

Focussing on the polychromatic erythroblasts where changes in proteome were more pronounced, we interrogated the set of proteins significantly up-regulated and significantly down-regulated in *HBB*^−/-^ cells, also using Gene Ontology and Pathway Enrichment Analysis toolkits WebGestalt^27,28^ and DAVID^29,30^ to identify overrepresented pathways. No down-regulated pathways reached significance, but multiple significantly upregulated pathways were identified, as detailed below.

β-thalassemia has been described as a protein-aggregation disorder^31^ with increased activity of the ubiquitin-proteasome pathway and autophagy^32–34^ as cells attempt to clear the α - globin aggregates. Here we show that the ubiquitin-proteasome and autophagy pathways are significantly upregulated in *HBB*^−/-^ polychromatic erythroblasts (Fig. 5B), with the majority of proteins identified in our dataset involved in these pathways increased in level (Fig. 6A and B). In addition, the abundance of the majority of proteins in the lysosome biogenesis pathway, associated with autophagy, was increased (Fig. 6C). We also found enrichment of pathways associated with heat shock response (Fig. 5B), with a striking increase in majority of identified proteins including many chaperones (Fig. 6D). Changes in abundance of proteins within these pathways were also detected in basophilic erythroblasts, although in general were less apparent than in polychromatic erythroblasts (Fig. 6), in line with increasing α -globin levels.

**Figure 5:**
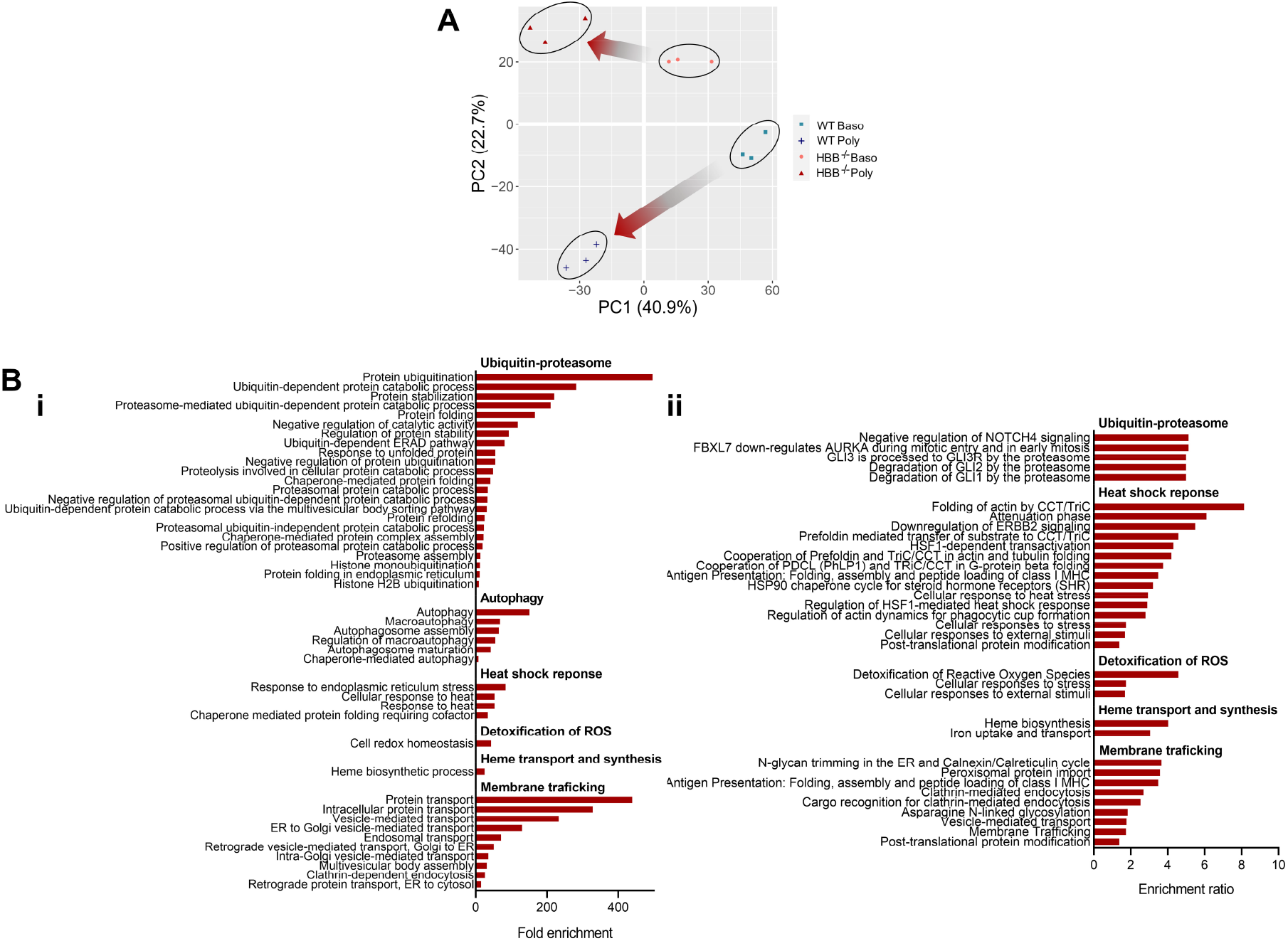
Comparative proteomic analysis of *HBB*^−/-^ and WT BEL-A. (A) Principal component analysis (PCA) plot of multiplex TMT-based comparative proteomics of FACS-isolated basophilic and polychromatic erythroblasts from *HBB*^−/-^ and WT BEL-A. (B) Overrepresented pathways in HBB^−/-^ vs WT BEL-A polychromatic cells from TMT-based comparative proteomic data. Pathways shown were significantly enriched by FDR (<.05) from (i) DAVID (ii) Webgestalt proteomics pathway analysis tools.

**Figure 6:**
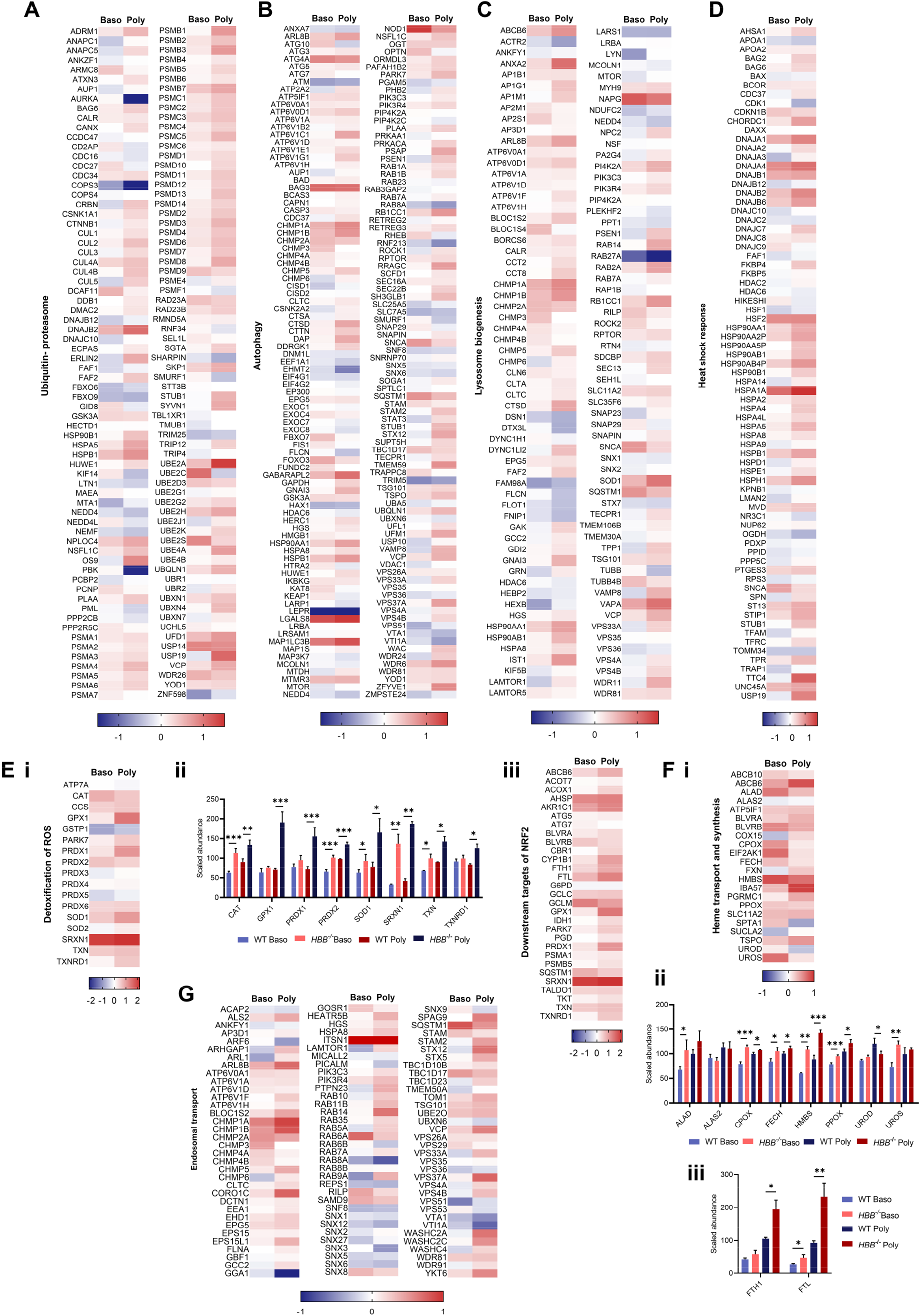
Analysis of alterations in abundance of proteins from significantly enriched pathways of *HBB*^−/-^ vs WT BEL-A. Heatmaps show log2 fold-changes (FC) of all WT vs *HBB*^−/-^ normalised protein abundances in both basophilic (Baso) and polychromatic (Poly) erythroblasts from significantly upregulated pathways identified by pathway analysis, using the following search terms on http://geneontology.org/: (A) ubiquitin-proteasome, (B) autophagy, (C) lysosome biogenesis, (D) heat shock response, (Ei) detoxification of reactive oxygen species (ROS), (F) heme transport and synthesis, and (G) endosomal transport. (Eii) Heat map of all downstream target proteins of NRF2 identified in the dataset. Bar charts show abundance values normalized to total protein and scaled relative to 100 shown as mean ± SD, n=3. \**P* < .05, \*\**P* < .01, \*\*\**P* < .001. *P* values represent results from ANOVA performed on log2 normalized data.

Of note, the most significantly expressed chaperone protein in both *HBB*^−/-^ polychromatic and basophilic erythroblasts was HSP70 (HSPA1A; Supplementary Figure 8). HSP70 has previously been reported to interact with aggregated α-globin in β-thalassemia erythroblasts sequestering it to the cytosol, thus preventing its translocation to the nucleus to protect GATA1 from degradation^5^. Consistent with this finding, GATA1 levels were significantly decreased in *HBB*^−/-^ basophilic and polychromatic erythroblasts (Supplementary Figure 8).

Few pathways involved in oxidative stress have been studied in detail in β-thalassemia erythroid cells^35,36^ although auto oxidation of α-globin aggregates is hypothesised to lead to increased ROS^37^. Consistent with this, pathways associated with detoxification of ROS were significantly overrepresented (Fig. 5B), with the majority of proteins involved in detoxification of ROS found at increased levels (Fig. 6Ei), including eight antioxidant enzymes significantly increased in polychromatic erythroblasts, with four also significantly increased in basophilic erythroblasts, prior to onset of apoptosis (Fig. 6Eii). In addition, target genes of NRF2^38–40^, an activator of the antioxidant response programme^41^ previously shown to be upregulated in β^IVS-2-654^ thalassaemic mice^40^, were increased (Fig. 6Eiii).

Interestingly, pathways involved in heme transport and synthesis were overrepresented in our dataset (Fig. 5B), with proteins involved in heme biosynthesis found at overall higher levels in both *HBB*^*-*/-^ basophilic and polychromatic erythroblasts (Fig. 6Fi). This included seven of the eight core heme synthesis enzymes, with just one in basophilic and two in polychromatic erythroblasts failing to reach significance (Fig. 6Fii). Levels of both ferritin light chain and ferritin heavy chain were also significantly increased (Fig. 6Fiii). In relation, pathways associated with endosomal transport and membrane trafficking were significantly enriched in polychromatic erythroblasts (Fig. 5B) and the majority of proteins involved in endosomal transport found at increased levels (Fig. 6G), indicating that more general transport mechanisms may be upregulated in order to elevate heme transport capacity.

Overall, these findings suggest increased heme biosynthesis in β-thalassemia erythroblasts, likely in response to decreased hemoglobin levels, which could contribute to erythroid intrinsic iron overload and thus potentially to the IE phenotype.

## Discussion

In this study we created, characterised, and validated human disease cellular model systems for β-thalassemia, which recapitulate the phenotype of patient erythroid cells. The lines provide a sustainable and consistent supply of disease cells as unique research tools for studying the underlying molecular defects, for identifying new therapeutic targets, and as screening platforms for novel drug and therapeutic reagents. For the latter we also developed a high throughput compatible fluorometric based assay for evaluation of severity IE and thus of disease phenotype. We then utilised the assay to demonstrate our β-thalassemia lines respond appropriately to drug and gene therapy approaches previously shown to increase γ-globin, with resultant improvement to IE, providing validation for such applications.

Proteomic studies have previously been utilised to identify potential biomarkers of β-thalassemia disease severity, analysing serum^42^, plasma^43–46^, or peripheral blood microvesicles^47,48^, for improved clinical management. Such analyses, although important, address downstream sequelae of the root cause of disease pathophysiology, IE of the erythroid cells. Our β-thalassemia disease lines overcome the hurdle of obtaining adequate numbers of bone marrow erythroid cells or haematopoietic stem cells from patients, enabling in depth and accurate analysis of molecular changes underlying the defective differentiation. In addition, an issue with comparing control and β-thalassemia erythroid cells arises from differences in kinetics of the cultures^3^ (Fig 1B and 3B), whereby differences due to stage of differentiation can confound identification of those due to disease phenotype. However, as we show, our lines enable sufficient size cultures to isolate specific, stage-matched cells.

Using TMT-based comparative proteomics we reveal the changes in biological pathways and processes in β-thalassemia erythroid cells, including upregulated pathways associated with, and increased levels of, antioxidant response enzymes. In addition, we identified increased heat shock proteins and chaperones, and increased levels of many proteins of the ubiquitin-proteasome and autophagy pathways along with proteins of associated lysosome biogenesis, together with respective upregulated pathways. The changes in proteome are manifest in earlier erythroid cells, but with the magnitude of change increased by the polychromatic erythroid cell stage, in line with increased levels of α -globin. However, this plethora of cellular responses are unable to adequately remove the aggregated α-globin or resolve the downstream sequelae, resulting in manifestation of the disease phenotype.

Previously, only few proteins associated with some of these processes have been reported upregulated in β-thalassemia erythroid cells^32,34,49^, although increased abundance of transcripts for most proteosome subunits has been reported in the β-globin^Th3/+^ and β-globin^Th3/Th3^ mouse ^33^, in line with increased levels of these proteins in human cells in the present study. In contrast, our data reveal extensive up regulation of proteins involved in wide range of biological response pathways, in a biologically relevant human system.

We also reveal upregulated pathways and increased levels of proteins for heme biosynthesis, including all eight core heme synthesis enzymes, along with associated upregulated endosome biogenesis and trafficking pathways. These findings support the hypothesis that erythroid-intrinsic iron overload may be contributing to oxidative stress in β-thalassemia, as heme molecules bind insoluble α-globin chains exacerbating the toxicity of these aggregates^50^. Heme also inhibits HRI, increasing globin synthesis, which will cause further aggregate formation^51^. Regulated reduction of components of heme synthesis in erythroid cells may therefore provide potential novel avenues for therapeutic intervention. In keeping, catalytic activity of erythroid specific ALAS2, the first enzyme in the heme synthesis pathway, can be modulated by small molecules binding a hotspot around its autoinhibitory Ct-extension^52^. Alternatively, use of succinylacetone, an inhibitor of the second step of heme synthesis, and the iron chelator Eltrombopag have shown promise in alleviating disease phenotype in Diamond Blackfan Anemia (DBA) erythroid cells *in vitro*^53,54^, further supporting the potential benefits of an iron reduction approach to treating β-thalassemia.

Intriguingly, we also found a novel mutant β-globin splice variant arising from the IVS-1-1 G→ T mutation, predicted to reduce the stability of dimer and tetramer and thus still resulting in a β^0^-thalassemia phenotype. A transcript for this variant has previously been reported from overexpressing IVS-1-1 G→ A *HBB* coding region construct in Hela cells^25^. That the variant does not have a beneficial role correlates with the lack of evidence for a milder clinical phenotype in IVS-1-1 G→ T patients compared to those with frameshift mutations^55^. In addition, there was no evidence of reduced insoluble α-globin in our IVS-1-1 compared to CD41/42 line, further supporting the hypothesis that the IVS-1-1 β-globin variant cannot form stable interactions with α-globin. Instead, as the variant β-globin precipitates it could exacerbate the disease phenotype. As there are many splicing mutations that cause β-thalassemia, it will be interesting to determine if other in-frame mutant β-globin splice variants are produced and if so, how they impact disease phenotype.

Finally, the findings presented here also provide proof of principle for CRISPR-Cas9 genome editing of BEL-A to create sustainable model cellular systems of other red blood cell disorders.

## Methods

### Cell culture

BEL-A cells were cultured as previously described^56^. In brief, for expansion phase cells were cultured in StemSpan™ SFEM [Stemcell Technologies] containing 50 ng.ml^−1^ SCF, 3 U.ml^−1^ EPO, 1 μM dexamethasone and 1 µg.ml^−1^ doxycycline. To induce differentiation, expanding cells were transferred to erythroid differentiation medium (Iscove’s medium with stable glutamine [Merck] containing 3% (v/v) AB serum [Merck], 2% (v/v) fetal bovine serum [Hyclone], 10 µg.ml^−1^ Insulin [Merck], Heparin 3U.ml^−1^ [Merck], 200 µg.ml^−1^ holo-transferrin [Sanquin, NED] and 3U.ml^−1^ EPO [Roche]) supplemented with 1 ng.ml^−1^ IL→ 3 and 10 ng.ml^−1^ SCF (both R&D Systems) and 1 µg.ml^−1^ doxycycline for 4 days, and for a further 4 days without doxycycline. Cells were then transferred to erythroid differentiation medium supplemented with holo-transferrin to a final concentration of 500 µg.ml^−1^. For hydroxyurea treatment, hydroxyurea (H8627; Sigma) was added to cells at 50 μM every 2 days from 4 days prior to the start of differentiation until harvesting of cells for analysis.

### CRISPR-Cas9 genome editing of BEL-A

#### Plasmid-based editing

To generate *HBB*^−/-^ and *HBB*^+/-^ lines, Guide RNA oligos (GTAACGGCAGACTTCTCCTC) were ligated into the pSpCas9(BB)-2A-GFP (px458) vector (48138; Addgene). BEL-A cells (1.0 × 10^6^) were resuspended in CD34+ nucleofection kit buffer (Lonza Biosciences) containing 2.5 µg px458 (with gRNA) and 2.5 µM ssODN donor template, and electroporated using program U-008 of AMAXA Nucleofector™ 2b. After 48 h, dead and dying cells were removed using a dead cell removal kit (130-090-101; Miltenyi). After a further 24 h, single-cell cloning of GFP+/DRAQ5-cells into 96-well plates was performed using a BDInflux Cell Sorter.

#### Ribonucleoprotein (RNP)-based editing

A sub-cloned population of BEL-A cells (1.0 × 10^5^) were resuspended in CD34+ nucleofection kit buffer (Lonza Biosciences) containing 18 pmol Cas9, 45 pmol gRNA. To generate the *HBB*^−/-^ *HBD*^−/-^ line, three gRNAs previously validated by Boontanrart et al.^57^ were used simultaneously, two of which were specific to HBB (e87 (g10) CUUGCCCCACAGGGCAGUAA; e383 GCUCAUGGCAAGAAAGUGCU), and one of which targeted both *HBB* and *HBD* (e297 UGGUCUACCCUUGGACCCAG). For CD41/42 -TCTT IVS1-1 G→ T *HBB* edits, gRNAs ((CD41/42 *HBB* - GGCUGCUGGUGGUCUACCCU; IVS1-1 *HBB* – AAGGUGAACGUGGAUGAAGU, Synthego) were used along with 100 pmol ssODN template containing the mutation of interest. ssODN templates were antisense for both the CD41/42 and IVS1-1 edits with homology arms 36 nucleotides away from the edit and 91 nucleotides towards the edit, in accordance with the design principles described previously^24,58^. A recoded template was chosen for IVS1-1 G→ T, where silent mutations are incorporated into the ssODN between the site of the DSB and the intended edit to increase editing efficiency^24^, due to the relatively large distance from the edit to DSB.

BCL11A +58 enhancer editing, was performed as above with HBB^−/-^ BEL-A as a founder population using a previously validated gRNA targeting the *BCL11A* +58 enhancer (CUAACAGUUGCUUUUAUCAC) ^2,13^.

#### Clone screening

Genomic DNA was isolated and the gene region of interest was amplified using GoTaq DNA polymerase (Promega) (*HBB* Fw: 5’-TGGTATGGGGCCAAGAGATA-3’, *HBB* Rv: 5’-GAGCCAGGCCATCACTAAAG-3’; *BCL11A +58* Fw: GGCAGCTAGACAGGACTTGG, *BCL11A +58* Rv: GGAGGCAAGTCAGTTGGGAA). PCR products were cleaned up (QIAquick PCR purification Kit, Qiagen) and sequenced by EuroFins Scientific. Sequencing data were analysed by TIDE^59^ and TIDER^60^ web tools (https://tide.nki.nl/).

#### Cytospin preparation, imaging, and morphology analysis

Aliquots of 2-8×10^4^ cells in 200 µl of appropriate cell culture medium were used to prepare cytospin preparations on coated slides, by centrifugation at 400 *g* for 5 min with a Thermo Scientific Shandon 4 Cytospin. The slides were stained in Leishman’s Eosin-Methylene blue solution (VWR). The stained slides were imaged using an Olympus CX43 microscope mounted with Olympus SC50 camera. Morphology analyses were determined by counting at least 200 cells per slide.

### Flow cytometry

#### Ineffective erythropoiesis (IE) assay

Aliquots of 2-3×10^5^ cells were incubated with CD36-Vioblue conjugated antibody (130-095-482; Miltenyi) in PBS containing 1% (w/v) BSA (Park Scientific Ltd) and 2 mg.ml^−1^ glucose (PBS-AG), followed by incubation with Annexin V-FITC (130-093-060; Miltenyi) or Annexin V-APC (640920; Biolegend) conjugated antibody in Annexin V binding buffer (10mM Hepes, pH7.4; 140mM NaCl; 2.5mM CaCl2). Cells were analysed on a BD LSR Fortessa flow cytometer and data analysed using FlowJo v10.6.1 (FlowJo LLC).

#### Intracellular staining of γ-globin

Following IE assay staining as above, cells were fixed in PBS-AG + 4% PFA + 0.0075% glutaraldehyde and permeabilised in PBS-AG with 0.1% saponin. Cells were blocked in in PBS containing 4% BSA and 0.1% saponin before incubation with HbF-FITC conjugate antibody (130-108-241; Miltenyi). Data collection and analysis was performed as above.

### RP-HPLC

RP-HPLC was performed as described by Loucari *et al*.^61^ with the exception of the injection size (600,000 cells in 30 µl dH2O) and the LC column used (Jupiter 5µm C18 300A, size 250 × 4.4 mm, protected with a Security Guard analytical guard system [KJ0-4282, Phenomenex]).

### SDS-PAGE and western blot

Proteins (either from whole cell lysates or insoluble fraction) were resolved by SDS-PAGE and transferred to PVDF membrane (Merck) by western blotting. The insoluble fraction was obtained by centrifugation (17,000 *g*) of whole cell lysate homogenised in RIPA buffer, before direct resuspension in SDS-PAGE sample buffer, with the equivalent of 250,000 cells loaded per lane. For whole cell lysates, 3 µg of protein was loaded per lane. Membranes were blocked with 5% (w/v) milk powder before incubation in primary and HRP-secondary antibodies (α-globin, sc-514378; β-globin, sc-21757; γ-globin, sc-21756 [Santa Cruz]; β-actin, A1978 [Merck]; BCL11A, ab19487 [Abcam]). Bands were visualized using enhanced chemiluminescence (G.E. Healthcare) with a G:BOX Chemiluminescence imager (Syngene).

### Cell stage matching for comparative proteomics

GPA (Glycophorin A) vs CD36 cell surface marker analysis by flow cytometry was used to isolate stage-matched basophilic and polychromatic erythroblasts for WT and *HBB*^−/-^ BEL-A cells. Cells were obtained from 3 independent cultures.

### TMT labelling, mass spectrometry and data analysis

TMT LC-MS/MS comparative proteomic experiments were performed as previously described^56^ with the following adaptions. Aliquots of 100 µg of protein from whole cell lysates were digested with trypsin (2.5 µg trypsin, 37 °C overnight) and labelled with Tandem Mass Tag (TMT) eleven-Plex reagents according to the manufacturer’s protocol (Thermo Fisher Scientific) before labelled samples were pooled.

Data analyses were performed on log2 normalised abundances using an un-paired, 2-tailed, heteroscedastic Student’s *t*-test in Microsoft Excel. A 5% false discovery rate (FDR) threshold for each comparison was calculated using R Studio. For data visualisation, principal component analysis (PCA) plots and volcano plots were generated using R Studio. Heatmaps were generated using GraphPad Prism 8. For overrepresentation analysis, significantly differentially expressed proteins (*P* < 0.05) were functionally categorized based on Reactome (https://reactome.org) via gene ontology (GO) classification using WebGestalt^27,28^ (http://www.webgestalt.org/). Gene ontology biological process was used as a functional database. Functional annotation of biological pathways was also performed using DAVID^29,30^ (https://david.ncifcrf.gov). WebGestalt and DAVID use the Benjamini-Hochberg Procedure to decrease the false discovery rate (FDR). Significantly enriched pathways are presented (FDR ≤ 0.05).

### Molecular modelling

The model for the IVS-1-1 mutant was built using Alphafold^62^. The 1VS-1-1 mutant model was then aligned with the β-globin proteins in the high-resolution X-ray structure of human hemoglobin (pdb code: 1A00^63^). The tetramer was relaxed by energy minimisation with the Gromacs software^64^ using the Gromos 54A7 force-field^65^. The system was energy minimized with the steepest-descent method in order to remove excessive strain by performing 100 steps of minimization with harmonic restraints applied to all non-hydrogen atoms, followed by further 100 steps restraining the Cα atoms only, ending with 100 steps with no restraints.

### Statistical analysis

For statistical analysis of >2 groups, following Shapiro-Wilk normality testing, ANOVA with Tukey multiple comparison testing was performed. For analysis of 2 groups, a Welch’s *t*-test was performed. In both cases, statistical tests were performed in GraphPad Prism 8.

## Acknowledgements

The authors would like to thank Dr Kate Heesom, Director of the University of Bristol Proteomics Facility, for performing mass spectrometry and analysis of raw data, Dr. Phil Lewis, School of Cellular and Molecular Medicine, University of Bristol, Bristol, UK for mass spectrometry data analysis, the University of Bristol Flow Cytometry Facility for use of equipment, and Dr Andrew Herman, director of, and Helen Rice of the Flow Cytometry Facility for performing cell sorting. The gRNAs used to generate the *HBB*^−/-^HBD^−/-^ cell line were a gift from Professor Jacob Corn, ETH Zurich, Switzerland.

The study was supported by MRC grant MR/S021140/1 and Wellcome Trust grant 209739/Z/17/Z. The work of JAS was supported by BBSRC-funded SWBio DTP. ASFO would like to thank EPSRC (grant number EP/M022609/1), BBSRC (grant number BB/R016445/1) and ERC (Advanced Grant PREDACTED https://cordis.europa.eu/project/id/101021207) for support.

## Contributions

JF conceived and supervised the study. Experiments were conceived and designed by JF and DED with contribution from IF-V and JH. *HBB*^−/-^ and *HBB*^+/-^ cell lines were generated by IF-V and DCJF. *HBB*^−/-^*HBD*^−/-^ cell line was generated by JH. *HBB*^−/-^ *BCL11A* +58 edited cell line was generated by EMF. CD41/42 and IVS-1-1 cell lines were generated by DED. DED conducted the majority of experiments, analysed data and prepared figures. TNA and EMF conducted experiments. MCW carried out the HPLC experiments and contributed to proteomics analysis. ASFO carried out molecular modelling with contribution from JAS. JF and DED wrote the manuscript.

