## Supplemental Figures for "Human cellular model systems of β-thalassemia enable in-depth analysis of disease phenotype"

**Supplementary Data**

**Supplementary Figure 1**

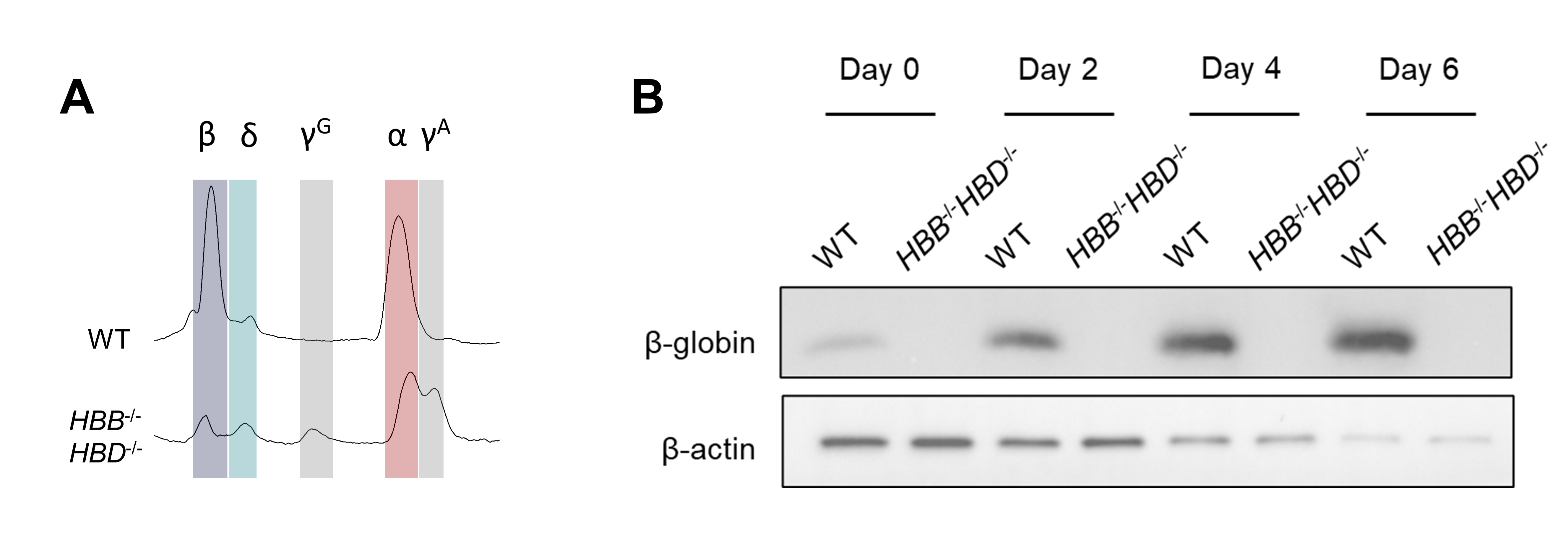

**Supplementary Figure 1: Confirming loss of β-globin despite residual peak on RP-HPLC overlapping position of β-globin.** These experiments were undertaken using a *HBB HBD* double knockout BEL-A line (*HBB*^-/-^*HBD*^-/-^) due to unavailability of a β-globin antibody that did not cross react with δ-globin. (A) RP-HPLC traces for WT and *HBB*^-/-^*HBD*^-/-^ BEL-A at day 6 of differentiation showing presence of small residual peaks in the β-globin and δ-globin positions. Normal peak positions are identified for β-globin (β), δ-globin (δ), ^G^γ-globin (^G^γ), α-globin (α) and ^A^γ-globin (^A^γ). (B) Western blot of whole cell lysates from WT and *HBB*^-/-^*HBD*^-/-^ BEL-A harvested at day 0, 2, 4 and 6 of differentiation, incubated with β-globin antibody, showing complete absence of β-globin in the *HBB*^-/-^*HBD*^-/-^ cells. β-actin was used as a protein loading control.

**Supplementary Figure 2**

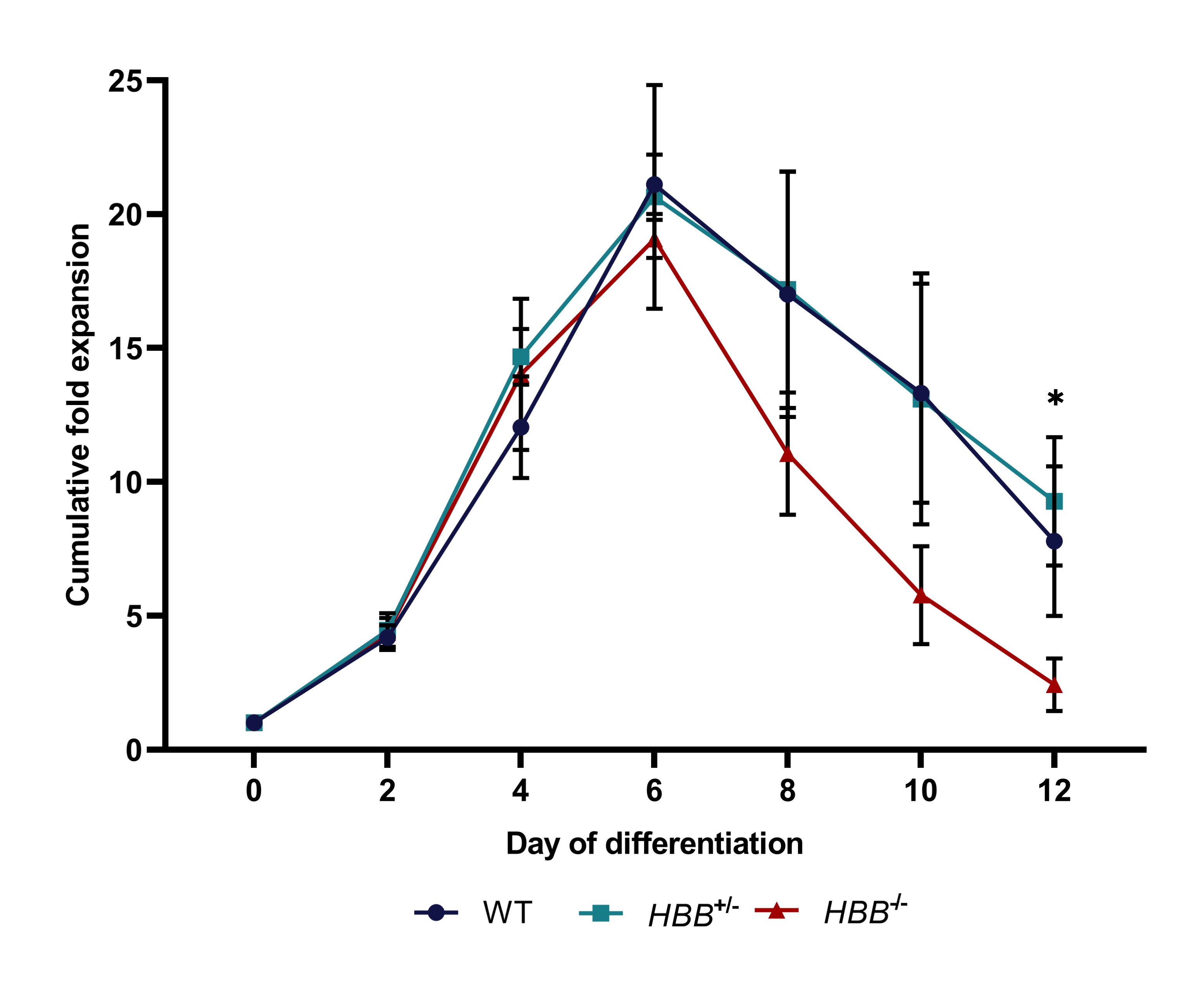

**Supplementary Figure 2: Expansion curves of WT, *HBB*^+/-^ and *HBB*^-/-^ BEL-A during erythroid differentiation.** Results show mean ± SD, n=3. **P* < .05.

**Supplementary Figure 3**

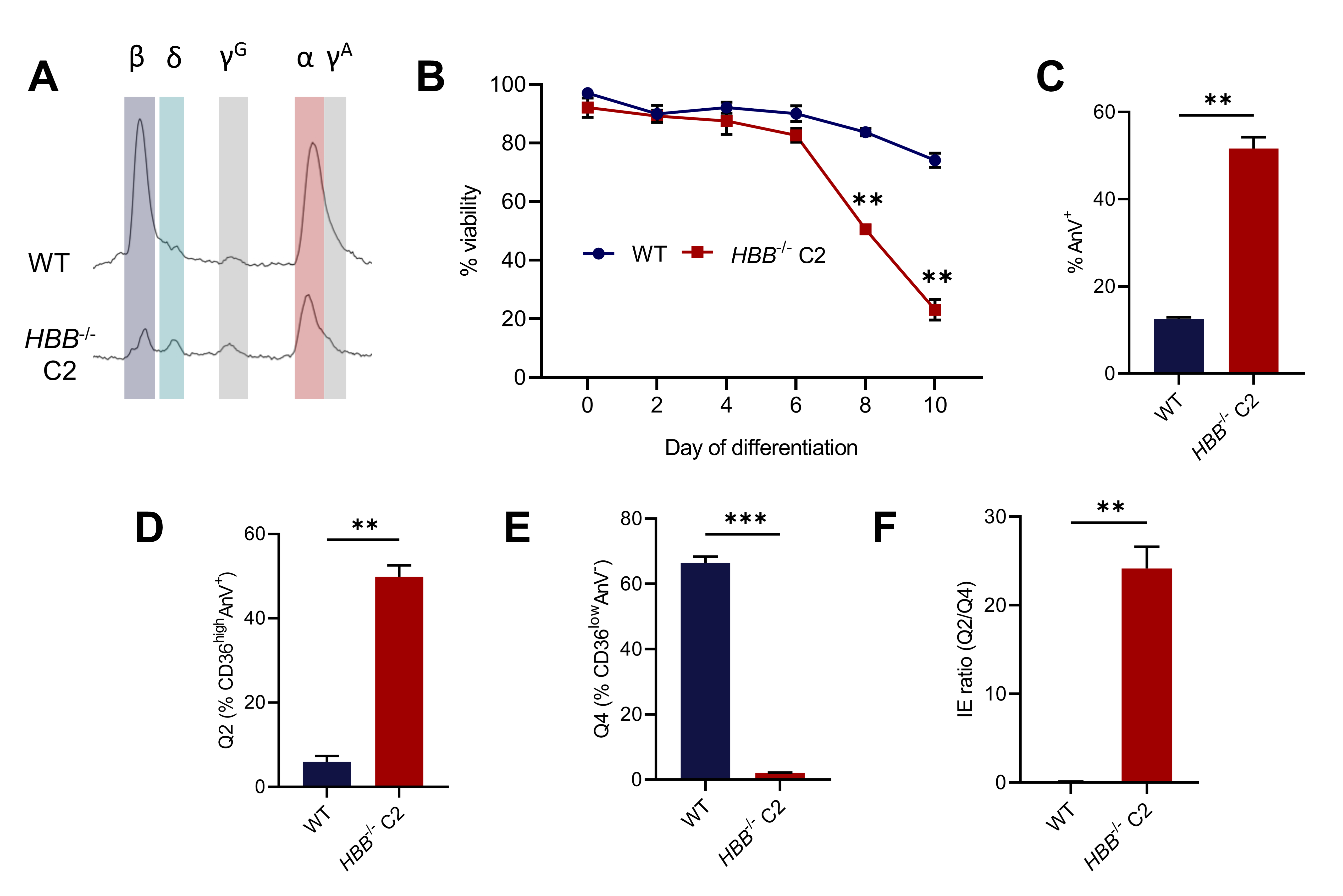

**Supplementary Figure 3: An additional clonal *HBB*^-/-^ BEL-A line demonstrates consistent phenotype**. (A) RP-HPLC traces for WT and *HBB*^-/-^ BEL-A at day 6 of differentiation. Peaks are identified for β-globin (β),  δ-globin (δ),  ^G^γ-globin (^G^γ), α-globin (α) and ^A^γ-globin (^A^γ). (B) Percentage viability during differentiation by trypan blue exclusion assay. (C) Percentage of Annexin V positive cells (AnV^+^) in at day 7 of differentiation as determined by flow cytometry. Quantification of Q2 CD36^high^AnV^+^ cells (D), Q4 CD36^low^AnV^-^ cells (E) and IE ratio (Q2/Q4) (F) at day 7 of differentiation. Results show mean ± SD, n=2. ***P* < .01, ****P* < .001.

**Supplementary Figure 4**

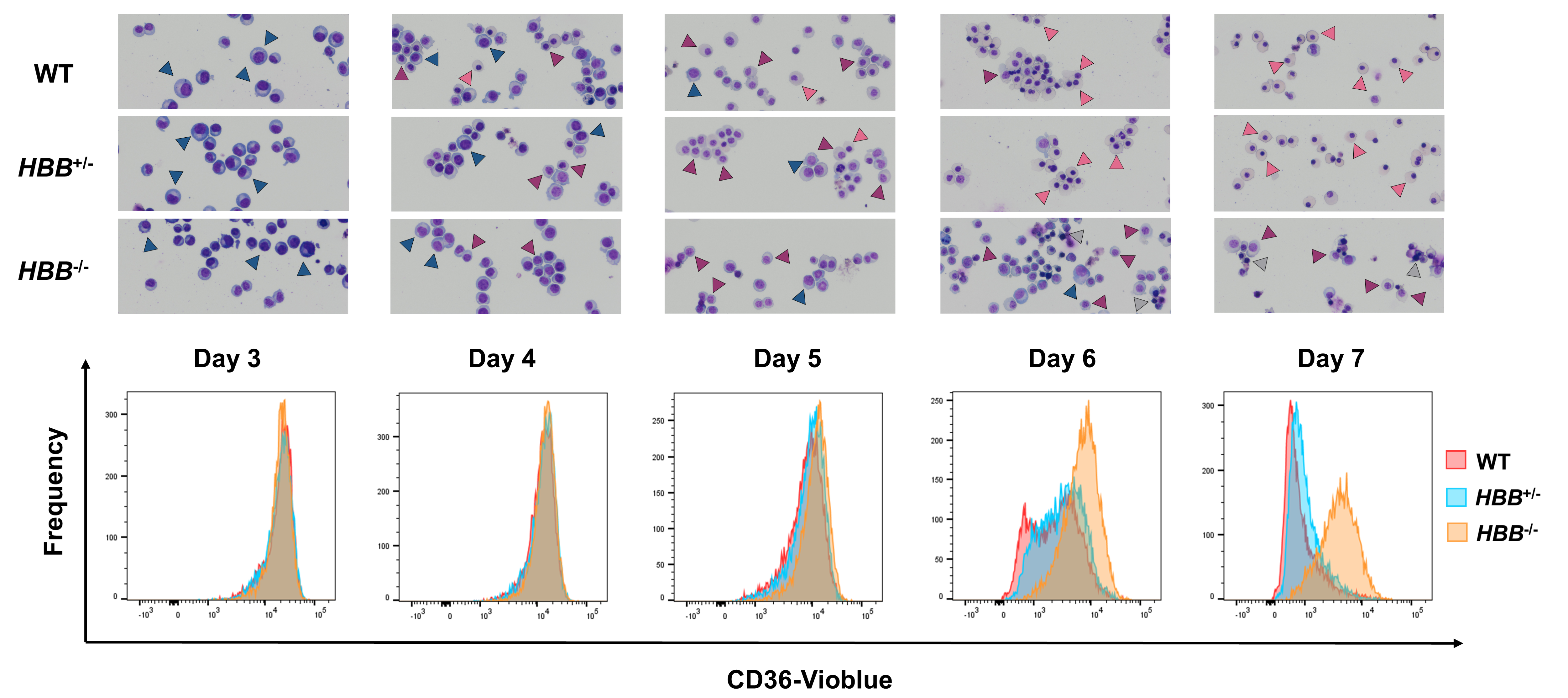

**Supplementary Figure 4: Flow cytometry analysis of CD36 cell-surface abundance in WT, *HBB*^+/-^ and *HBB*^-/-^ BEL-A cells with corresponding cytospin images during erythroid differentiation.** Arrowheads indicate the following cell types: blue, basophilic erythroblast; purple, polychromatic erythroblast; pink, orthochromatic erythroblast; grey, dead/apoptotic.

**Supplementary Figure 5**

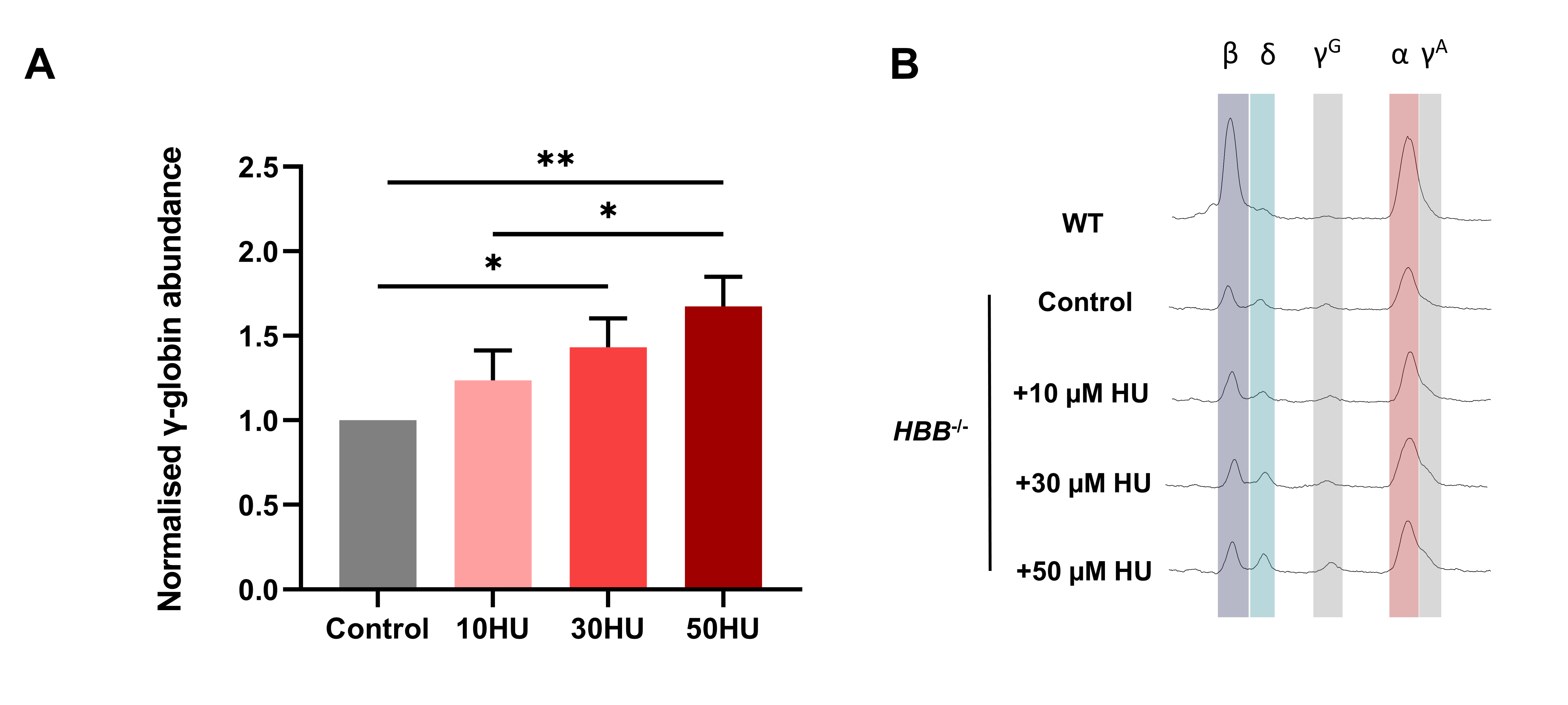

**Supplementary Figure 5: Gamma globin abundance in hydroxyurea (HU) treated HBB^-/-^ BEL-A.** Differentiating HBB^-/-^ BEL-A cells were treated with 10, 30 or 50 µM of HU and harvested for RP-HPLC analysis. (A) Abundance of total $\gamma$-globin (^G^γ-globin + ^A^γ-globin) at day 6 of differentiation quantified from RP-HPLC traces and normalised to total globin, shown as a proportion of control normalised $\gamma$-globin abundance. Results show mean ± SD, n=3. **P* < .05, ***P* < .01. (B). Representative RP-HPLC traces. Peaks are identified for β-globin (β), δ-globin (δ), ^G^γ-globin (^G^γ), α-globin (α) and ^A^γ-globin (^A^γ).

**Supplementary Figure 6**

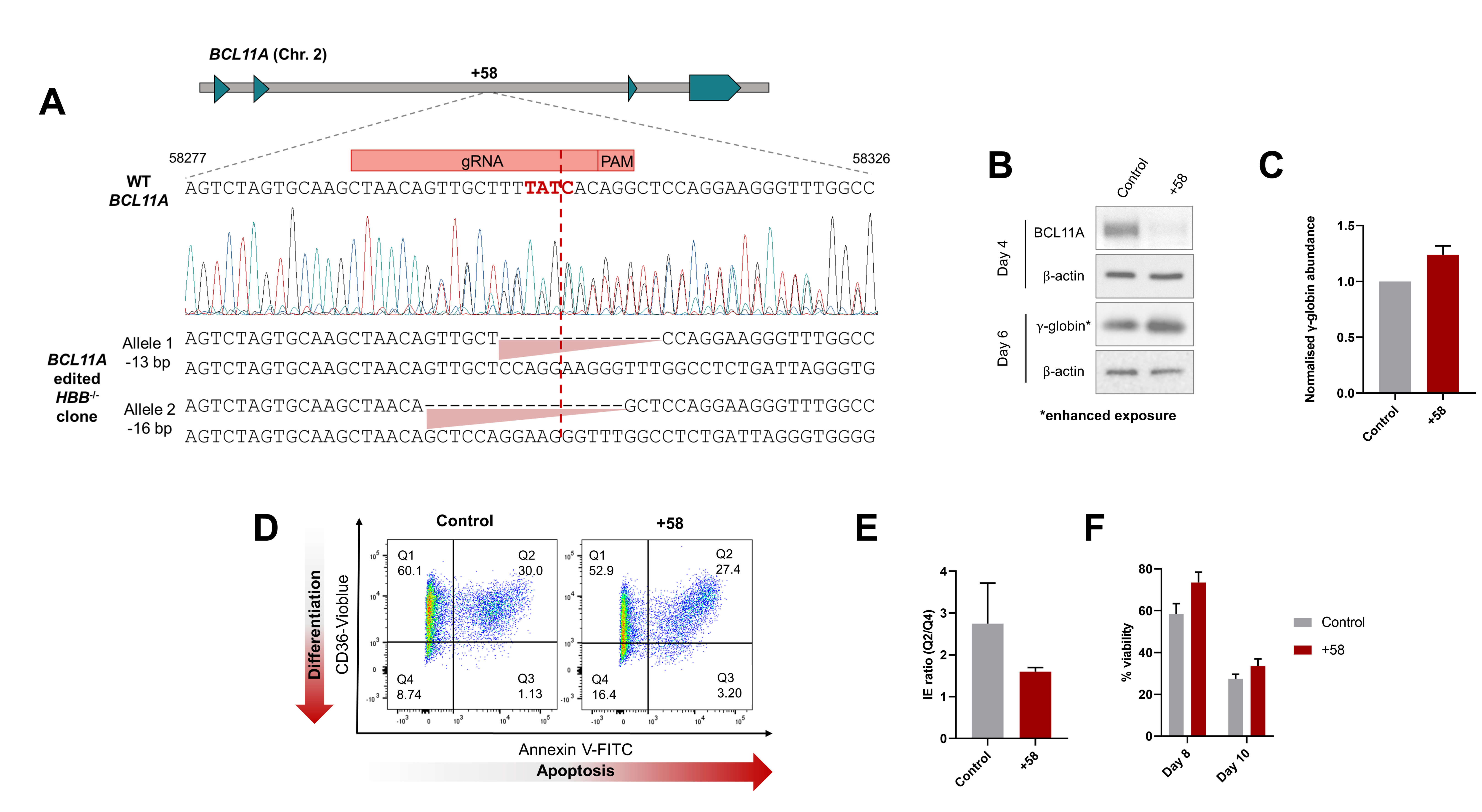

**Supplementary Figure 6: CRISPR-Cas9 genome editing of the *BCL11A* +58 enhancer in *HBB*^-/-^ BEL-A.** (A) Schematic showing gDNA sequence analysis of *BCL11A* +58 enhancer edited HBB^-/-^ BEL-A. Position of GATA box shown in red. (B) Representative western blots of control and +58 *BCL11A* enhancer edited *HBB*^-/-^ BEL-A cell lysates harvested at day 4 and 6, incubated with BCL11A and γ-globin antibodies respectively. β-actin was used as a protein loading control (C) Abundance of total $\gamma$-globin (^G^γ-globin + ^A^γ-globin) at day 6 of differentiation quantified from RP-HPLC traces and normalised to total globin, shown as a proportion of control normalised $\gamma$-globin abundance. (D) Representative IE flow cytometry plots of control and +58 *BCL11A* enhancer edited *HBB*^-/-^ BEL-A cells at day 7 and (E) Quantification of IE ratio (%CD36^high^AnV^+^/ CD36^low^AnV^-^). (F) Percentage viability of control and +58 *BCL11A* enhancer edited *HBB*^-/-^ BEL-A during differentiation by trypan blue exclusion assay. Results show mean ± SD, n=2.

**Supplementary Figure 7**

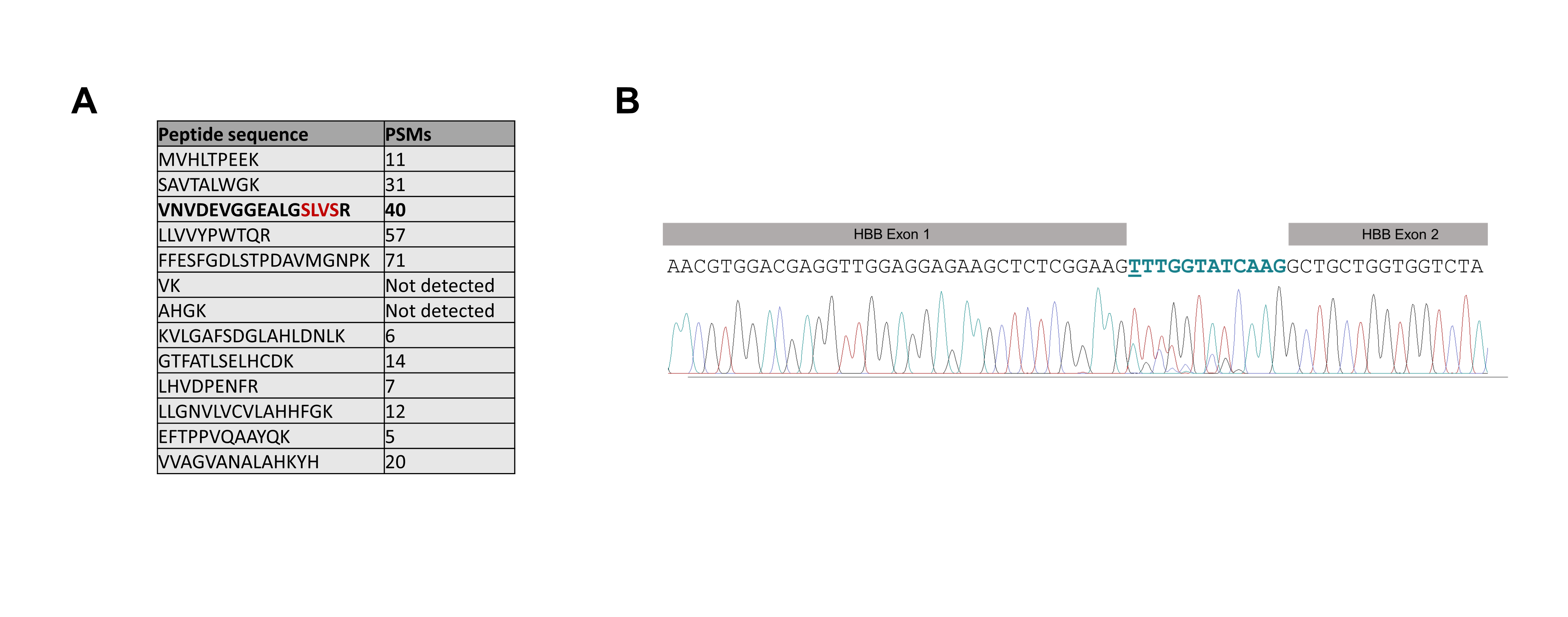
**Supplementary Figure 7: Identification of IVS-1-1 β-globin splice variant.** (A) Mass spectrometry results positively identify the predicted IVS-1-1 β-globin unique peptide sequence (bold) containing the additional 4 amino acids (red) from trypsin digest of unknown peak identified by RP-HPLC in Figure 3B. The number of peptide spectrum matches (PSMs) for each peptide from the trypsin digest of the variant β-globin are shown. (B) Chromatogram of IVS-1-1 BEL-A *HBB* transcript cDNA. Silent mutations resulting from the CRISPR edit are shown in red and the inserted 12 bases from the first intron of HBB as a result of splicing using the IVS-1-13 alternative splice site are shown in teal (with the G→T IVS-1-1 substitution underlined).

**Supplementary Figure 8**

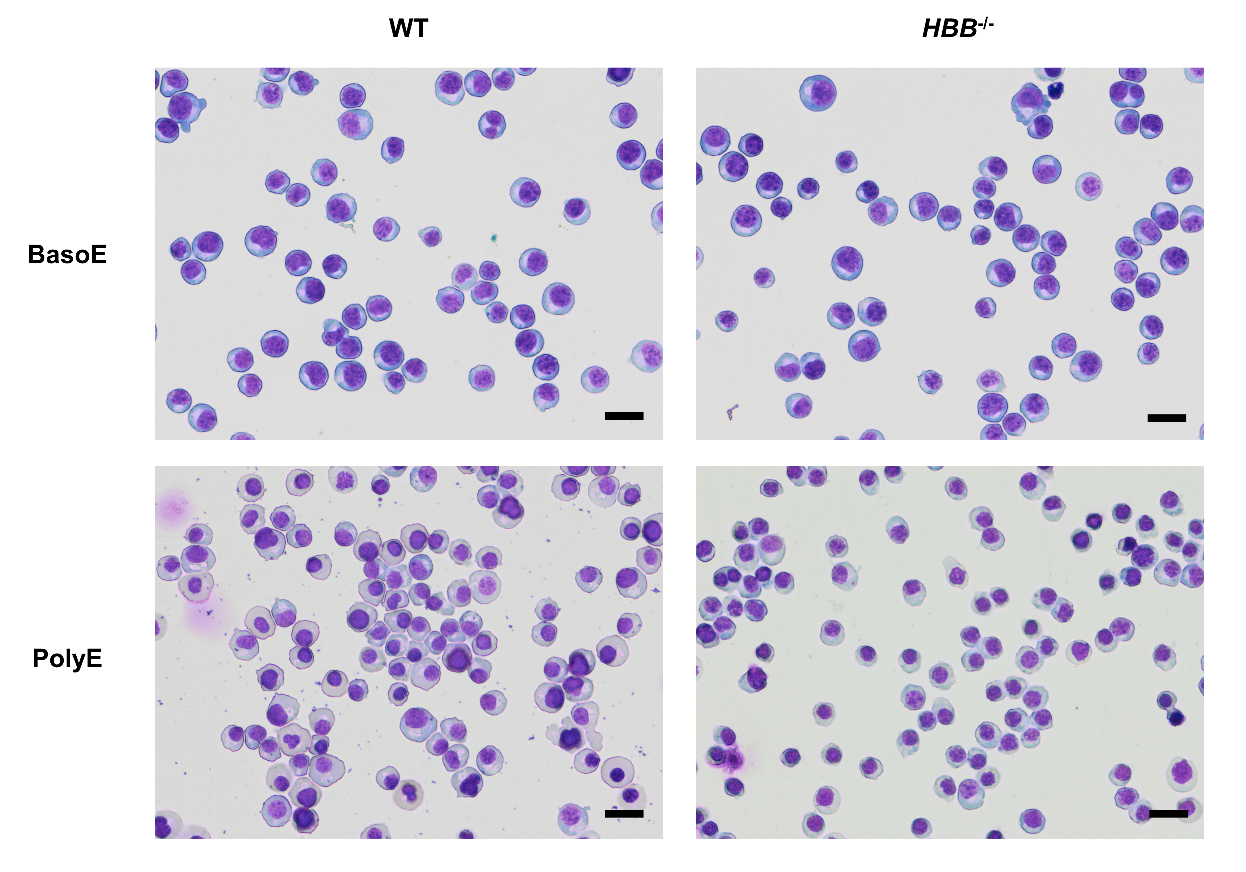

**Supplementary Figure 8: FACS isolated WT and *HBB*^-/-^ cells for multiplex TMT-based comparative proteomics.** Representative Leishman’s stained cytospin images of basophilic (BasoE) and polychromatic (PolyE) erythroblasts after FACS isolation based on cell surface expression of CD36 and GPA. Scale bars 20 μm.

**Supplementary Figure 9**

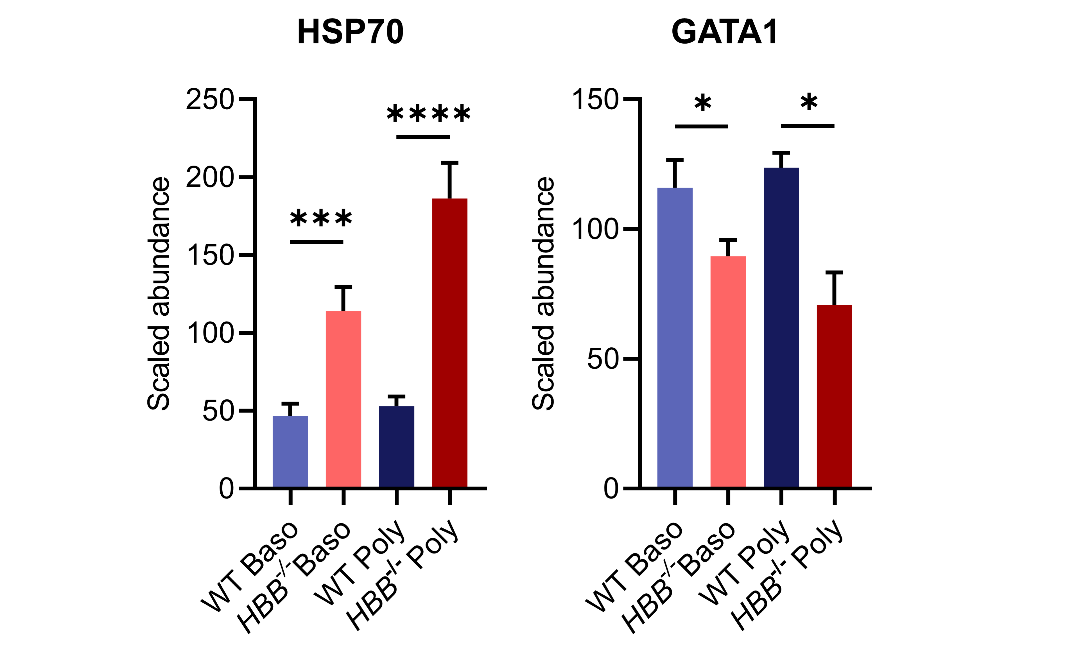

**Supplementary Figure 9: Scaled protein abundance of HSP70 and GATA1 in WT and HBB^-/-^ BEL-A cells from TMT-based comparative proteomic data .** Data shown are abundance values normalized to total protein and scaled relative to 100 shown as mean ± SD, n=3. ∗p < 0.05, ∗∗∗p < 0.001, ∗∗∗∗p < 0.0001. p values represent results from ANOVA performed on log_2_ normalized data.
